# Extending empirical dynamic modeling to cross-sectional data beyond traditional time series

**DOI:** 10.1101/2025.01.10.628762

**Authors:** Ethan R. Deyle, Gerald Pao, George Sugihara

## Abstract

The foundation of Empirical dynamic modeling (EDM) is in representing time-series data as the trajectory of a dynamic system in a multidimensional state space rather than as a collection of traces of individual variables changing through time. Takens’s theorem provides a rigorous basis for adopting this state-space view of time-series data even from just a single time series, but there is considerable additional value to building out a state space with explicit covariates. Multivariate EDM case studies to-date, however, generally rely on building up understanding first from univariate to multivariate and use lag-coordinate embeddings for critical steps along the path of analysis. Here, we propose an alternative set of steps for multivariate EDM analysis when the traditional roadmap is not practicable. The general approach borrows ideas of random data projection from compressed sensing, but additional justification is described within the framework of Takens’s theorem. We then detail algorithms that implement this alternative method and validate through application to simulated model data. The model demonstrations are constructed to explicitly demonstrate the possibility for this approach to extend EDM application from time-series trajectories to effectively realizations of the underlying vector field, i.e. data sets that measure change over time with very short formal time series but are otherwise “big” in terms of number of variables and samples.

## 1 Introduction

Time-series analysis with empirical dynamic modeling (EDM) has become increasingly visible as a means of addressing causality (1) and prediction in natural systems from scales of single neurons (2) to the entire Earth climate system (3). However, the continued growth in use cases and applications is necessarily limited by the availability of suitable time series. The general rule of thumb has been given that at least 30 time points are needed (4). This presents a challenge; in field, clinical and in vitro biological study, when sequential repeated measurements (time series) are made in large studies they are often quite short but then replicated across many individual systems, samples, or individuals. Increasingly, measurements are also coordinated across a huge breadth of different variables.

At present, the general approach to applying EDM to a new data set begins with univariate attractor reconstruction (5) and cross-map between univariate reconstructions (1) even when explicit (mechanistic) multivariate EDM models are ultimately derived and studied (6).

These traditional analyses generally require many sequential observations because the changes between sequential time points stand in for the unknown or at least yet-to-be determined interacting variables. Combining short, replicate time series (7,8) and using Multiview embeddings (9) have both been demonstrated as methods to enable EDM analysis on data sets that might otherwise not be of sufficient length.

Nevertheless, these extensions of the methodology don’t yet bridge to the most common case of emerging Big Data: the largest datasets from cells to coastlines are often cross-sectional. Their size and power come from repeating measurements across huge batches of cells, human subjects, or spatial locations with no repeated temporal measurement. Ultimately, truly cross-sectional data sets are extremely hard to use to explicitly study change-over-time and dynamics. But what if there are even just a few repeated measurements? Indeed, even if measurements are just repeated a single time for each instance (i.e. cell, human subject, spatial location), those data can still contain empirical records of the dynamics in a way purely cross-sectional data sets do not (see illustration Fig. 1). Imagine, for example, replicating the classic *Paramecium-Didinium* predator prey experiments (10) but instead of initializing all the cultures to have uniform starting conditions the initial concentrations of the ciliates were varied on an array. Re-sampling each culture after a short time 𝜏 (appropriate to the time scale of dynamics) can yield the same if not richer understanding of the dynamic patterns as allowing a single culture to evolve over an extended time course as in Fig 8 of (10).

**Fig 1.**
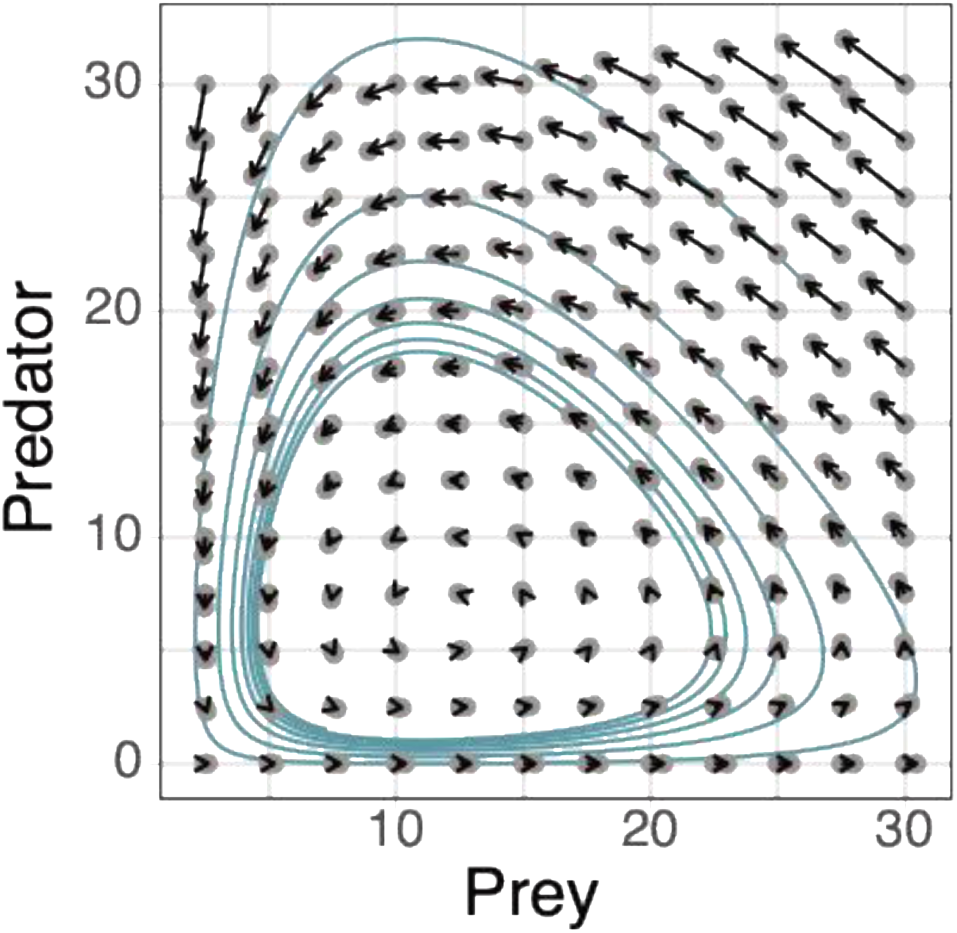
Schematic of dynamic data in classic predator-prey system. Traditionally EDM has been implemented with long unbroken time series that create continuous trajectories in state space (teal line). However, the reconstructing the vector field underlying the trajectories that evolve on top of it provides an alternative way to study the dynamic system without requiring traditional time series. Here we will show how the EDM approach can be applied to data sets consisting of many short “threads”. The most extreme example of this is shown as pairs of grey points connected by arrows, where many replicate systems are sequentially measured at just two time points, *t* and *t* + *𝜏*.

Of course, in this greatly simplified case there are just two variables and so creating this empirical vector field from the data is trivial. We can use existing EDM machinery like multivariate S-map to predict *Predator*(*t* + *𝜏*) or *Prey*(*t* + *𝜏*) as a function of *Predator*(*t*) and *Prey*(*t*). In real cases, there will be many possible candidate variables whether it’s 120 different species of reef fish or 6000 different gene sequences. If we could somehow know which sets of these variables already belonged together, we could simply proceed with multivariate EDM analysis. This is the original “trick” from Takens’s theorem that has powered empirical dynamic modeling. Time lags of a single variable serve as proxy variables for the dimensions of the dynamics that have yet to be determined or possibly even measured. In typical EDM implementations, this then enables further sequential modeling steps— (1) testing for the existence of attractor dynamics, using convergent cross mapping to test from hypothesized interacting variables which actually show evidence of shared attractor dynamics, and then (2) finally replacing those “proxy” coordinates to build out an explicit multivariate model. Once the final step is reached, there may no longer be any need to use the time-series structure of the data. For example, it is possible to create explicit multivariate embeddings that predict dinoflagellate blooms (11) from the available parallel measurements without using any time-lag coordinates.

Nevertheless, the machinery typically used to get there (univariate simplex projection, CCM, etc.) did in fact require the time-series structure.

Consequently, extending multivariate EDM analysis to cross-sectional data sets boils down to finding an alternative path to getting to final fully multivariate step. Previous multivariate methods of causal inference with EDM in ecological studies (12–14) taken together with insights into EDM from neuroscience applications (15) point a way forward. Rather than using univariate time lag coordinates as proxies to stand in for the yet-to-be determined interactors, we propose using proxy coordinates constructed from available co-variates using random linear combinations, i.e. random projection coordinates. Intuitively this lets us side-step the issue of which possible variable combinations are most directly causally related without being forced back to studying the system first through isolated pairwise correlations.

More formally, the foundation for this proposed approach is the multivariate generalization of Takens’s theorem (13), which makes explicit that a dynamic attractor can be reconstructed diffeomorphically from many various combinations of lags of different variables. This result highlights that with multiple interacting variables there is no single uniquely correct representation of a dynamical system. Furthermore, it immediately follows that any combination of simple transformations like coordinate rotations or other linear combinations of observation functions are also valid embedding variables. Thus, at least mathematically, there are an infinite number of valid ways to reconstruct a system from multiple time series. While this muddies the notion of which is the “best” set of variables to study a system, it also motivates multi-modeling approaches to multivariate empirical dynamic forecasting (9,16). Combining the predictions of multiple embeddings can smooth out forecast skill across areas of the attractor, improve forecast skill, and reduce uncertainty. When it comes to using EDM to quantify interaction strengths (12), however, the interpretations are most straight forward when a single set of multivariate coordinates can be selected and validated. Partial derivatives, after all, are embedding dependent. That is “how *x* changes with respect to *y* when all other state variables are held constant” depends very much on what other variables are being held constant (see Fig S1).

Below, we describe and investigate sequential analysis for multivariate EDM that exploits the multiplicity of valid embeddings while addressing the ambiguity that simultaneously arises.

Again, the essential notion is that randomly constructed combinations of possible interacting variables replace lagged time-series values as stand-in proxy coordinates through the intermediate EDM steps. We demonstrate through model investigations how the approach is a path for multivariate EDM analysis of data sets that contain some measurements of change over time without being classically constituted time series (see Fig 1 illustration). At the same time, the method can be of further use in standardizing the steps for choosing specific multivariate EDM models to focus deeper mechanistic investigations, ala (12). Thus, we also compare the success of the approach to heuristic algorithms applied in previous studies and investigate different implementations of the general approach for using random embeddings. Reassuringly, the particulars of the algorithm do not seem critical to the immediate usefulness of the general concept.

## 2 Methods

### 2.1 Multivariate approaches

#### 2.1.1 Multiview embedding

Multiview embedding (MVE) represents a simple but powerful approach to address the multiplicity of valid multivariate representations of coupled system dynamics (9). It is an extension of weighted nearest neighbor forecasting with simplex projection, invoking an ensemble of embedding models. In the original implementation, instead of nearest neighbor ***x****_n_*_(i)_ getting weighting based on its distance from the forecast target ***x***\* = ***x***(t*) on a single manifold as in simplex, the weighting in MVE is assigned based on how many multivariate models ***x****_n_*_(i)_ is the single nearest neighbor of ***x***\*. In practice, it is optimal for forecasting to only consider a limited number *k* of the top multivariate models ranked on forecast skill in a test set and a good heuristic is to take 𝑘 = √𝑚 where *m* is the total number of multivariate models considered. Chang et al. recognized the further possibility of connecting high dimensional data sets to S-map estimation of the interaction Jacobian proceeding from the multiview principle (16) of multivariate EDM ensembles. The basic idea involves a two-step process for generating S-map models between a large number of possibly interacting species and shows how ensemble embedding analysis can be a path towards robust multivariate EDM analysis.

#### 2.1.2 Multivariate EDM with Randomized Embeddings

Work in neuroscience by Tajima and collaborators (Tajima et al. 2015) introduced the idea of using random projection coordinates popular in other branches of machine learning and compressed sensing for empirical dynamic modeling. Tajima et al. used randomly generated linear combinations of univariate time lags to generate coordinates for EDM with the reasoning that these would be more robust to multiple timescales than strict, sequential lags. Mechanically, this procedure is really just applying a (generically) full rank linear transformation to the univariate reconstructed attractor which we are already assured is diffeomorphic to the “true” attractor, guaranteeing that (at least theoretically) the attractor reconstructed from these randomly selected combinations (projections) is itself diffeomorphic as well.

This basic idea can be immediately generalized to multivariate coordinates, and (as we demonstrate here) this insight provides solutions for applying EDM to high spatial, low temporal power data. Indeed, the result is essentially a lemma of the proof of mixed lag embeddings (13). If you have *N* observation functions {*Y_1_, Y_2_, … Y_N_*} that embed a system—which can be any arbitrary mix of lags of different variables ala (13)—then generically a linear combination of the *Y_i_* will also be a valid embedding coordinate. Thus, we can generate at random any arbitrary number of projection coordinates ξ_j_

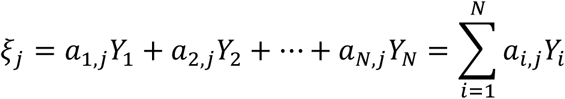

where the *a_i,j_* are scalar values drawn at random from the same distribution. Tajima et al. (15) note that Gaussian and Unitary distributions are both reasonable choices. If we replace a coordinate *Y_i_* with one of these random projection coordinates ξ_j_, the new manifold **M’** will be diffeomorphic to the old manifold **M**, because the linear transformation between the two sets of coordinates is non-degenerate for almost all random draws of a_i,j_:

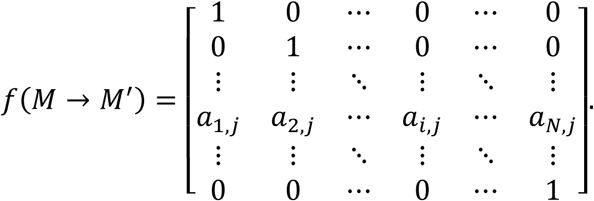

In practice, due to uncertain causal coupling between variables under consideration it will not necessarily be true that all variables being considered will in fact be valid embedding coordinates. In this case, including non-interacting variables in the random project coordinates contributes noise, sensu “estimation error” in (17). EDM forecasting methods like simplex and S-map are robust to noise up to a point, so the approach can be practical so long as the uncertainty added from including spurious variables does not degrade the predictive signal to the point of non-detection.

These randomized embeddings do not necessarily enhance predictive skill but rather can instead aid inference in system identification and causality detection. In particular, they allow a first attempt to reconstruct trajectories before a suitable embedding of causal variables is identified whether or not the data permit traditional univariate lag-coordinate embeddings. Note as well that there is a natural connection between these random multivariate embeddings and the previously mentioned multiview attractor reconstruction (9) in that the individual embeddings involve projection via sparse matrices (uniform coordinate sampling without replacement), and obey the same basic mathematics (18).

### 2.2 Proposed method

Building on these principles, we propose a methodology for constructing explicit multivariate EDM representations of system dynamics (embeddings) without using time-lag coordinates. In principle, the first step is to determine a rough estimation of the necessary embedding dimension, E. This can also be done with random projection coordinates in direct generalization of the original implementation of univariate random projection coordinates by Tajima et al. (15). However, the results of the model demonstrations presented below demonstrate that an exact estimation of embedding dimension isn’t critical. Thus, we focus on the additional steps, writing the algorithms using an *a priori* assumed embedding dimension, *E*\*.

The approach is outlined as follows. First, we extend the premise of cross mapping from a univariate lag coordinate attractor to a candidate time series (1), but now using an embedding space reconstructed with randomized coordinates. This calculation can then be applied as a test, i.e. if multivariate cross-map with candidate *Y_i_* predicts target *Y_j_* better than a surrogate, this is indicative of *Y_j_* having a causal influence on *Y_i_*. Next, just as with a traditional approach we sequentially replace the proxy coordinates with mechanistic variables to build out a mechanistic embedding from based on forecast improvement (14,19) from the subset of study variables that show evidence from multivariate cross-map of coupling. Finally, sequential application of the test can be used in a “greedy” approach to create a complete method for building an explicit multivariate state space.

#### 2.2.1 Multivariate Cross Mapping

As in traditional convergent cross mapping (CCM), the forecast target is not included in the embedding and the cross-map forecasting skill from the embedding coordinates to the target is used as an indicator of common membership to a low-dimensional dynamic system. This set up will allow a purely static treatment of the data where there are no temporal relationships among repeated observations. If there is a large initial set of candidate variables, this phase of analysis can be used to trim the list of possible embedding variables toward the more likely causal ones.

Given that, pseudo-code for the general algorithm is as follows. Given a target variable *Y*_1_ and *n*-1 other observational variables of the same system state {*Y*_2_, *Y*_3_, …, *Y*_n-1_} (*E*\* ≤ *n*-1):

i. Normalize all variables to have *mean*(*Y_i_*) = 0 and *sd*(*Y_i_*) = 1.
ii. Construct *E*\* random projection coordinates from the *n*-1 observed variables, and renormalize the ξ_j_

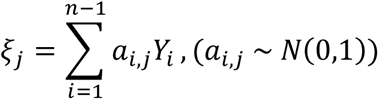

(iii) Compute the simplex projection forecast skill (see **Error! Reference source not found.**) of *Y_1_* for the embeddings including one candidate *Y_i_*, and *E*\*-1 random projection coordinates: {***Y_i_***, ξ_1_, ξ_2_, …, ξ_Emax-1_},
(iv) Compute a reference simplex projection forecast skill (see **Error! Reference source not found.**) of *Y_1_* for the embeddings using one of the following schemes:
a. A surrogate time series generated from the candidate *Y_i_*, and *E*\*-1 random projection coordinates: {∼***Y_i_***, ξ_1_, ξ_2_, …, ξ_Emax-1_},
b. Just the *E*\*-1 random projection coordinates: {ξ_1_, ξ_2_, …, ξ_Emax-1_},
(v) Repeat (ii)-(iii) for many ensembles (e.g. 500) randomly generated coordinates ξ_j_.

These calculations can be readily implemented using the *Simplex* function in either ‘rEDM’ (20) or ‘pyEDM’ (21). To increase statistical inference, the same ensemble of random projection coordinates can be used for each candidate embedding variable and so a paired test can be used. Note that some of the candidate *Y_i_* can be time-lags of each other if the data set permit, but if all *Y_i_* are simultaneous measurements than there need not be any temporal relationship between observations in the data set.

#### 2.2.2 Pairwise multivariate forecast analysis

Temporal forecasting can also be employed in the process of variable identification, especially if there are distinct hypotheses to test. In this case, the forecast target can be included in the embedding, and then the forecast skill of the target variable compared between similar embeddings can untangle information about external (e.g. environmental) drivers (19,22). The idea is that if *X_1_* and *X_2_* are both purported drivers of *Y_1_*, then whichever gives better prediction skill when used in embeddings to predict *Y_1_* is a more proximate driver. This can be tricky for comparing just a single set of embeddings, however, e.g. a univariate embedding of *Y_1_* with just a single lag of one of the drivers, {*X_1_*(t), *Y_1_*(t), *Y_1_* (t -τ), …, Y_1_(t – (*E*–2)τ)} and {*X_2_*(t), *Y_1_*(t), *Y_1_* (t - τ), …, Y_1_(t – (*E*–2)τ)}, and again cannot be readily applied to cases where using many time-lag coordinates isn’t possible.

Instead, this can be made more rigorous and robust using random projection coordinates and following a similar algorithm to that described previously for determining embedding dimension with random coordinates, but instead of comparing embedding with different numbers of randomized coordinates, we compare {*X_1_*, *Y_1_*, ξ_1_, ξ_2_,…, ξ_E-1_} and { *X_2_*, *Y_1_*, ξ_1_, ξ_2_,… ξ_E-1_}.

#### 2.2.3 Greedy EDM model selection with random projection coordinates

Following from this pairwise principle of forecast improvement, past EDM work with long time series has built multivariate models by replacing univariate lag coordinates with explicit variables (11). A similar approach can be used with randomized coordinates. This is motivated by two considerations. First, it is especially practical if the data do not allow for multiple time lags, e.g. data sets with many replicates but very short time series (see again Fig 1 illustration). In this sense it can be a technique to build predictive multivariate EDM models while circumventing the standard univariate analysis to determine embedding dimension (23). The second advantage is that for each observed variable, it is possible to generate a whole distribution of forecast skill at each stage of variable selection, and thus deciding on the single best variable at each stage can be done much more robustly because it is done with a multi-model approach.

The proposed algorithm is largely the same as for Multivariate Cross Mapping above, merely differing in the treatment of the target variable. Again, we assume an approximate embedding dimension *E\** has already been determined. Given a target variable *Y*_1_ and *n*-1 other observational variables of the same system {*Y*_2_, *Y*_3_, …, *Y*_n-1_}, some of which may be time-lags of each other, pseudo-code for the general algorithm is as follows:

i. Normalize all variables to have *mean*(*Y_i_*) = 0 and *sd*(*Y_i_*) = 1.
ii. Construct *E_max_* - 2 random projection coordinates from the *n*-1 observed variables, and renormalize the ξ_j_

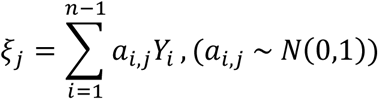

(iii) Compute the simplex projection forecast skill (see **Error! Reference source not found.**) of *Y_1_*(t + tp) for the embeddings including *Y_1_*, one candidate *Y_i_*, and *E*\*-2 random projection coordinates: {*Y_1_*, ***Y_2_***, ξ_1_, ξ_2_, …, ξ_Emax-1_},{*Y_1_*, ***Y_3_***, ξ_1_, ξ_2_, …, ξ_Emax-1_}, …, {*Y_1_*, ***Y_n_***, ξ_1_, ξ_2_, …, ξ_Emax-1_}. (Bolding used for emphasis).
(iv) Repeat (ii)-(iii) for many ensembles (e.g. 500) randomly generated coordinates ξ_j_.
(v) Select *Y_i_* that gives the highest median forecast skill.
(vi) Repeat (ii)-(v) but replacing a random projection coordinate with the *Y_i_*, and removing *Y_i_* from the candidate variables.

To increase statistical inference, the same ensemble of random projection coordinates should be used for each candidate embedding variable.

### 2.3 Models

Two toy models are used to demonstrate the proposed approach. The first is the multispecies competition model described by Huisman and Weissing for their Fig 1 (25). The second model is constructed using a coupled set of Repressilator circuits (26). We describe this gene circuit model in greater detail below, and code to reproduce the exact simulations presented in this manuscript is included in the supplemental material.

The basic Repressilator circuit first demonstrated by Elowitz and Leibler is shown in Fig 2A largely following the variable and parameter symbols later used by Volkov and Zhurov (27).

**Fig 2.**
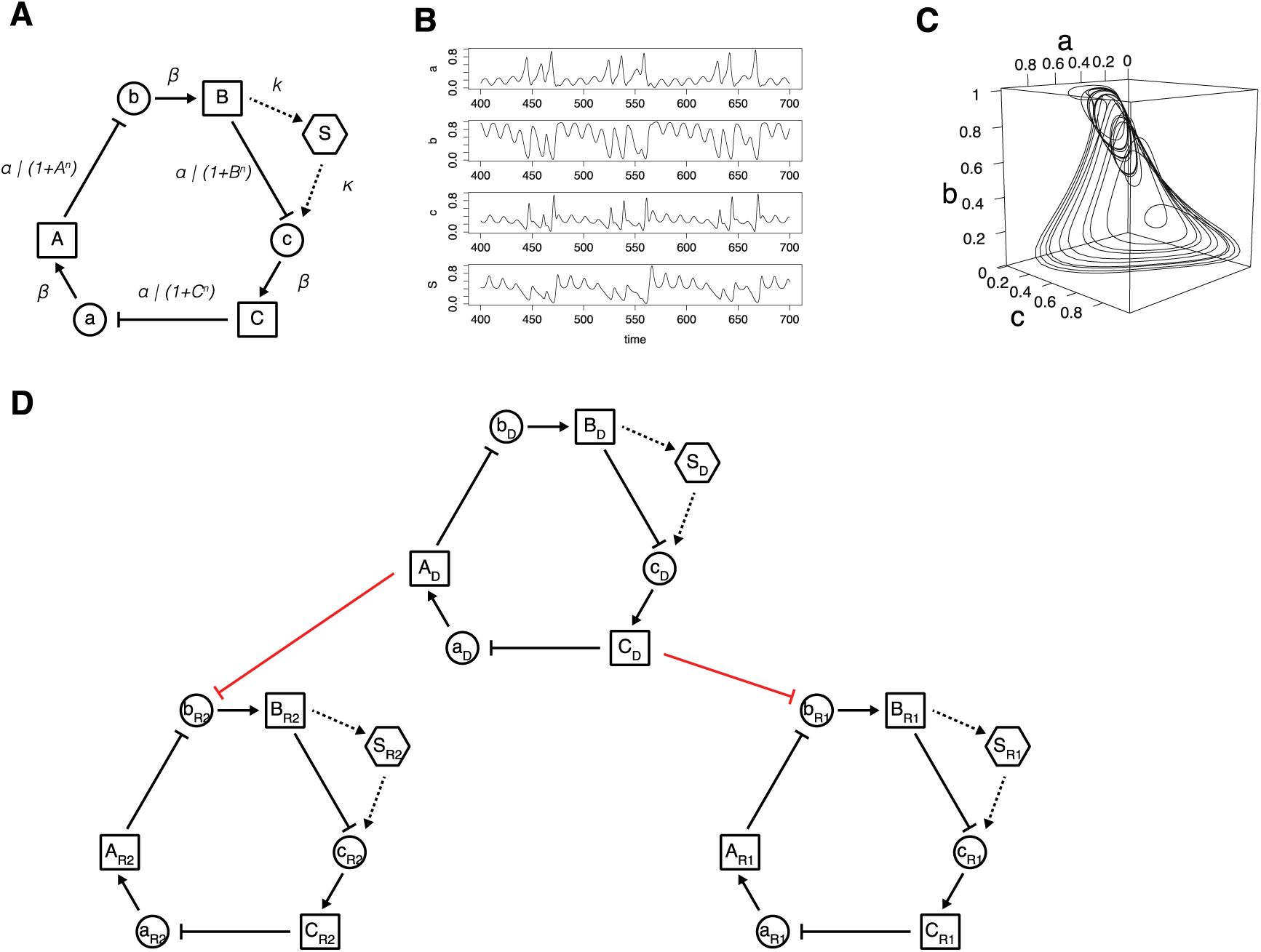
Networked repressilator model. A single Repressilator circuit is (panel A) can be tuned to produce quasiperiodic chaotic oscillations in expression levels (panel B) that follow a low-dimensional attractor manifold (panel C). A larger model is constructed by replicating the basic circuit three times, then adding a repressing interaction in one direction from one driving circuit to each of the other two response circuits (panel D). There are 23 parameters in total; each circuit has 7 parameters *α*, *β*, *κ*, *ks0*, *ks1*, *h*, and *η*, and there is a final parameter for each response circuit γ controlling the strength of the repressing action of the driver. Note that the respective parameter values are adjusted across circuits to avoid synchrony; these are given in Table S1.

It is composed of a rock-paper-scissors relationship between three genes, where gene *a*’s product *A* represses gene *b* expression, *b*’s product *B* represses gene *c* expression, and *c*’s product *C* represses gene *a* expression. Under certain parameter values, this produces quasi-periodic chaotic oscillations. Example time-series outputs are shown in Fig 2B, and the corresponding attractor view in Fig 2C.

This motif can then be used to construct more elaborate causal network that have low dimensional dynamical behavior that is mirrored in natural biological systems (28). We construct a network of 9 genes with a driving circuit that is unidirectionally coupled to two response circuits shown in Fig 2D. The driving Repressilator circuit with genes *a*_D_, *b*_D_, *c*_D_ is coupled to the first response circuit with genes *a*_R1_, *b*_R1_, *c*_R1_ by the product of the driver *A*_D_ repressing expression of response gene *b*_R1_. The driver product *C*_D_ represses expression of *b*_R2_. Thus, the causal effects between genes in the driving circuit and genes in R1 and R2 are unidirectional, while the genes in R1 and R2 do not have a causal interaction with each other but share common drivers (i.e. “case 3” sensu (1)) and do not exhibit synchrony under the parameterizations we use. We simulate additional time series variables that have no dynamical relationship whatsoever by sampling trajectories from simulations with different initializations.

## 3 Results and Discussion

### 3.1 State-space Identification with Randomized Coordinates

We apply the algorithm for Multivariate Cross Mapping with randomized coordinates as described above to a set of data sampled at random time points from a long realization of the Repressilator network model. Thus, all the data samples represent states of a dynamic system, but all information on sequence (trajectories) has been destroyed. This data set is thus an idealized representation of strictly latitudinal (single snapshot) laboratory or field measurements of many independent but equivalent systems or samples. Beginning with Fig 3, we show the change in forecast error (*ΔNRMSE*) under four alternative implementations of the general algorithm. We test both randomly selected multivariate coordinates and randomly projected multivariate coordinates; in each case a given embedding with the candidate coordinate is compared to either the *E*-1 dimensional embedding with the candidate removed or the *E* dimensional embedding with the candidate replaced with a randomized surrogate. Further explorations are then presented in supplemental figures varying some of the meta-parameters of the numerical experiment.

**Fig 3.**
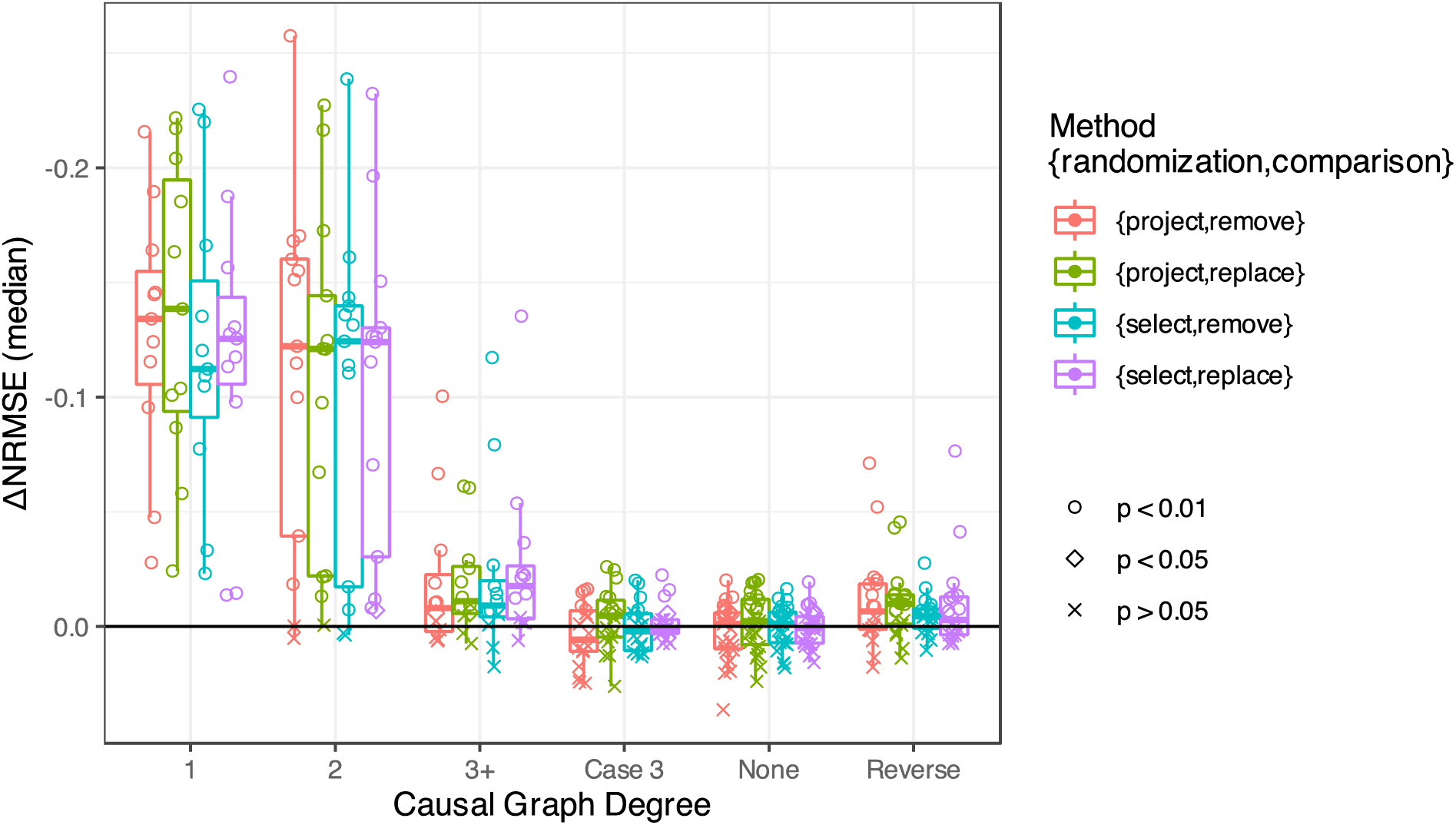
Forecast skill improvement under different implementations of the general multivariate test based on the degree of causal coupling between variables in the Repressilator network (*N_r_* = 100, *L* = 200, T:S = 1/4, and *E* = 6). The median difference in normalized root-mean squared error (ΔNRMSE) across the 100 coordinate re-randomizations is shown for each pair of target and candidate model variables. The specific implementations of the multivariate test algorithm are indicated by color. The shape reflects the p-value of a paired Wilcoxon test on the distribution of the mean squared errors between the test and reference (either removing or replacing) under re-randomization. The pairs are categorized by their true causal relationships described by the degree of separation between the target variable “i” and candidate variable “j” on the directed causal graph. Boxplots are drawn to summarize among the pairs within each category.

The results are grouped into distributions according to the pattern of their true underlying interaction. A pair that has a direct true causal interaction has a degree of “1”, a pair that have an indirect causal interaction mediated by another variable a degree of “2”, and indirect mediated by multiple other variables “3+”. Pairs that do not interact but share a mutual driver are categorized as “Case 3”, pairs that have a true interaction but only in the reverse direction to the test are categorized as “Reverse”, and pairs that have none of these associations but are from fully independent realizations of model systems are categorized as “None”. For the 9 genes in the Repressilator network model there are 81 possible directed pairs spread across all these categories aside from “None”. Finally, three spurious time series are included that are generated from an independent model realization of a single Repressilator are included to give 18 more pairwise tests with the causal degree “None”.

A preliminary comparison is shown in Fig 3, restricting possible numerical parameters of the test to specific values: *N_r_* = 100, *L* = 200, T:S = 1/4, and *E* = 6. Regardless of the specific way the general multivariate test is implemented, it is generally very clear to differentiate drivers that have a close interaction (degree of “1” or “2”) from reverse causal associations (“reverse”), shared drivers (“Case 3”), or no coupling relationship of any kind (“none”). In this initial experiment, there does not appear to be a clear advantage in using random projection coordinates (“project”) over random selected coordinates (“select”). Since we are repeating the test with multiple (*N_r_* = 100) different randomization of the coordinates, we are generating a distribution of the test statistic (ΔNRMSE) and therefore ascribe some certainty or significance to the difference in this measure of forecast skill with and without the candidate causal variable using a paired Wilcoxon test. These are also indicated on Fig 3 by point shape.

A wider exploration is shown in Fig S2 and Fig S3, where the number of extraneous variables stays fixed at T:S = 1/4, the embedding dimension *E* used for EDM prediction is varied from *E* = 3 to *E* = 9, and the number of vectors in the library between *L* = 50 and *L* = 500. For clarity, the test statistic ΔNRMSE is shown first as simple grouped box-and-whisker plots (Fig S2) and the corresponding p-values (taking the p-value of the Wilcoxon test as a proxy for a p-value of the EDM test) are visualized separately in Fig S3 as frequency histograms.

We can see some patterns in the types of errors made when the system is under- or over- embedded. When the system is over-embedded (*E* = 9), the procedure is more likely to incorrectly indicate a causal interaction where no causal relationship exists whatsoever.

However, when the system is under-embedded (*E* = 3), the procedure is more likely to miss a potentially meaningful causal interaction (e.g. degree = 2) than when over-embedded (*E* = 9). These patterns appear to hold whether there is a relatively small sample of data (*L* = 50) or not (*L* = 500). If we look at differences in the relative performances between the different particular methods within these broader patterns, however, it is hard to identify clear pattern and consequently to identify a clear “winner”.

Fig S4 shows a similar exploration to Fig S3 but keeping the number of training vectors constant (*L* = 200) and instead varying the ratio of true interacting variables to spurious (T:S = 1/2, 1/4, and 1/32). Although it might feel counter-intuitive, there appears to be some advantages of including spurious embedding variables in some ways. Particularly when the system is over- embedded (*E* = 3), the rates of false significance in “Case 3”, “Reverse”, and “None” are considerably lower. This is evident across the four different variations of the general methods, but the differences appear more pronounced using randomly projected embedding coordinates than randomly selected embedding coordinates. The purpose in exploring changes in T:S ratio was inspired by thinking of different use cases where the number of possible interacting variables could be quite large, but the possible benefit could be further explored. Intentionally adding spurious embedding variables does resemble the use of “noise injection” to regularize neural network training (29) and further investigation might show the mechanism of the effect is itself similar.

Additionally, we reflect on if the significance based on simply comparing distributions of prediction skill is a good indication of the significance of the cross-map inference of causality.

These p-values are only heuristically being ascribed to the causal test; they are in fact the p-value of a different statistic. Examining Fig S4 for example shows that a stricter threshold on the Wilcoxon p-value (p < 0.001) largely differentiates lack of interaction from interaction cases, so while this p-value itself is not directly meaningful taken literally, there is a clear suggestion that a more appropriate p-value could be arrived at with a more conservative test. Thus, more careful approaches, e.g. with surrogate methods, could sharpen the test. On the other hand, we are often interested in identifying causal variables for the purpose of modeling to produce forecasts with mechanistic interpretability. We do not need an exhaustive catalogue of all the true interactors, but instead an approximate embedding of the necessary dynamics so that we attain a near maximal forecast skill and can have confidence in estimates of pairwise interaction motifs (12). For this mode of study, it is therefore more important that direct drivers show a higher test statistic than indirect drivers or confounded variables, not that the test significance is perfectly refined.

Finally, as noted above, there is not a universal clear advantage either way between using randomly selected multivariate coordinates or randomly projected multivariate coordinates in these model simulations. Recall that for all the above results the number of random trials per pairwise test is large (*N_r_* = 100). However, basic intuition pulled from compressed sensing is that random projection coordinates as implemented here should be more “efficient” in a sense than random coordinates selected by sparse projection. Thus, we perform the same trial as shown in Fig 3, but using only 20 random draws for each pairwise trial. Fig 4 shows that indeed the random projection approach appreciably reduces the variance in test statistic (here ΔNRMSE) for each variable pair when only 20 random draws are used, demonstrating that the clearest advantage of the random projection approach is that it can be deployed with greater computational efficiently than random coordinate selection. We additionally consider the effect of changing the number of vectors in the training library (*L*), and the robustness largely holds even limiting the training library to *L* = 50.

**Fig 4.**
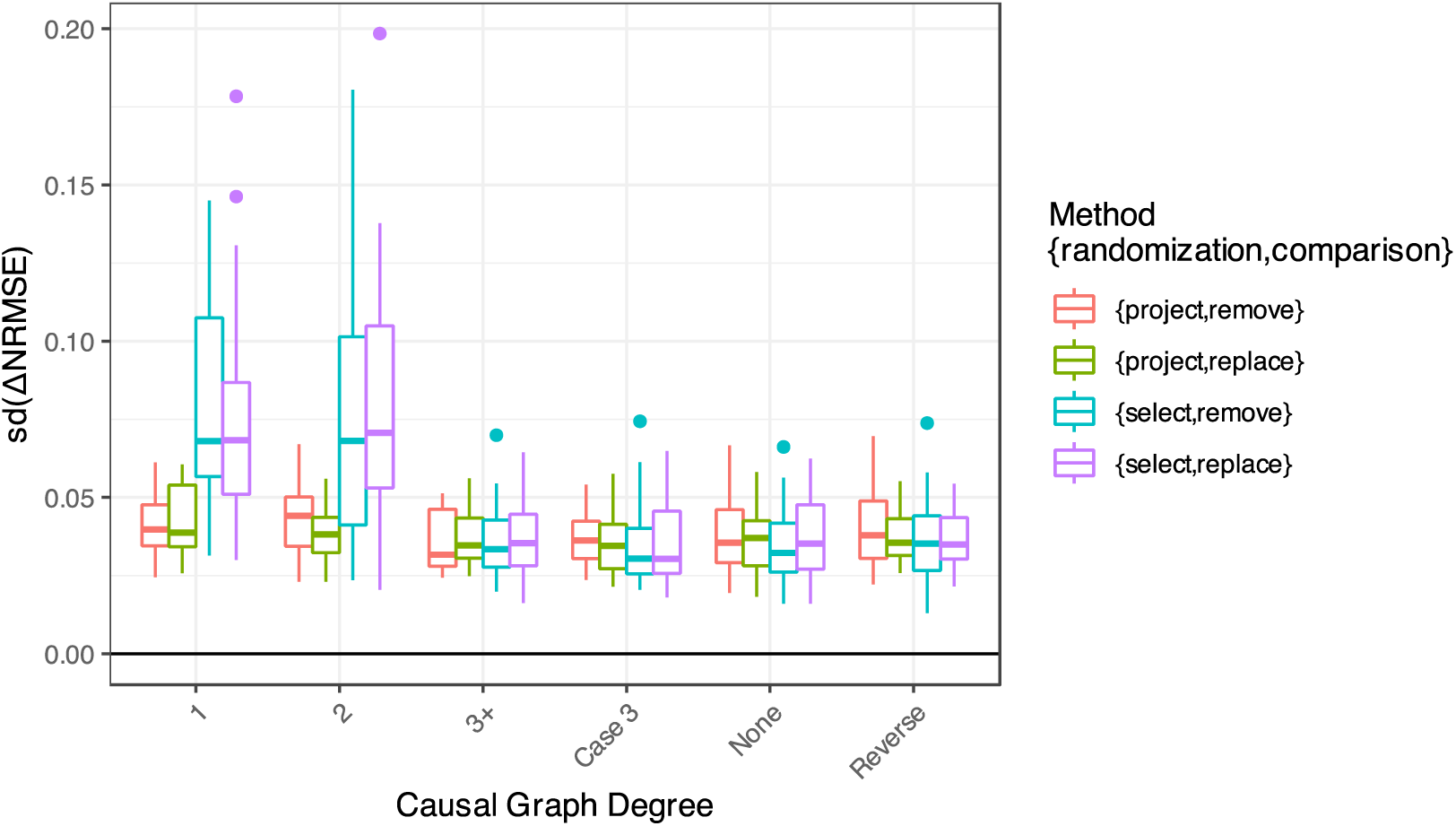
Efficiency of random coordinate approaches at small random sample size (*N_r_* = 20, *L* = 200, T:S = 1:1, and *E* = 6). Calculating the standard deviation of the forecast improvement (-ΔNRMSE) under a smaller number of re-randomizations (*N_r_* = 20) shows that using randomly projected coordinates rather than randomly selected coordinates can converge on a robust estimate more quickly.

### 3.2 Greedy Search with Randomized Coordinates

If this multivariate approach can correctly identify the most proximal interactors, it should be possible to string the test together to identify a complete set of coordinates that embed the system. The notion of what constitutes a complete embedding, or even more restrictively the *best* embedding is tricky to pin down in nonlinear biological systems. These model systems have a dissipative nature matching our general regard for the real systems— the dimension of the attractor manifold itself is lower than the full order of the system of differential equations generating them. The Huisman multispecies competition model is a straightforward example: the model is posed with *m* species (5) competing for *k* resources (3) but the dynamics can be understood entirely (i.e. the system can be completely embedded) with just the *m* population abundances. Consequently, it is a bit of a tricky issue to even say for certain what the correct “full embedding” is; and in fact, this really isn’t of concern when thinking about the central use- case for EDM application— to real data without a knowable true answer. Therefore, we center testing on out-of-sample forecast skill as this is the gold standard we have for observational data science. We compare a greedy search building up a candidate embedding by sequentially replacing randomized coordinates with the best performing candidate to using a simpler greedy search without random coordinates (in the vein of (6)) and to exhaustive search over combinations (more in the vein of (11)).

In nearly every trial, a simple greedy search without random coordinates yields a multivariate model with significantly lower out-of-sample forecast skill than either exhaustive search or random projection. Comparing exhaustive search and random projection further, there does appear to be some measurable danger of overfitting when using an exhaustive search for embedding coordinates when the time series are sufficiently long (*N* = 500), visible e.g. in the lower quantile of the red boxes being lower than green or turquoise in all but one case in the left-hand panels. This is directly examined in Fig S6 by looking specifically at the rate of including at least one erroneous variable in the course of model selection. We see in the short data series case (*L* = 50), exhaustive search and random projection are largely similar in the rate of overfitting (including erroneous variables), but that for longer series (*N* = 500) the error rate is noticeably higher for exhaustive search. At the same time, the simple greedy search is substantially less prone to identifying erroneous variables, and thus the lower out-of-sample forecast skill is best understood as arising from including suboptimal coordinates or under-embedding (not including enough coordinate variables).

Looking across these results (Fig 5 and S6), the performance is often very similar between exhaustive search and greedy search with random coordinates, with both approaches selecting embeddings in-sample that have similar forecast skill when then tested out-of-sample. However, these calculations are not trivial for current computing architecture. The number of individual simplex projection calculations that need to be run the full set of possible multivariate coordinates gives 15,504 unique model combinations (20 coordinates chosen 5 at a time). This number can quickly become unmanageably large for more candidate coordinates and large embedding dimension. For example, if we allowed an additional time lag for each species variable and an embedding dimension of 6, then we would need to contend with 593,775 multivariate forecasts. The greedy search depends on the number of random coordinate trials.

**Fig 5.**
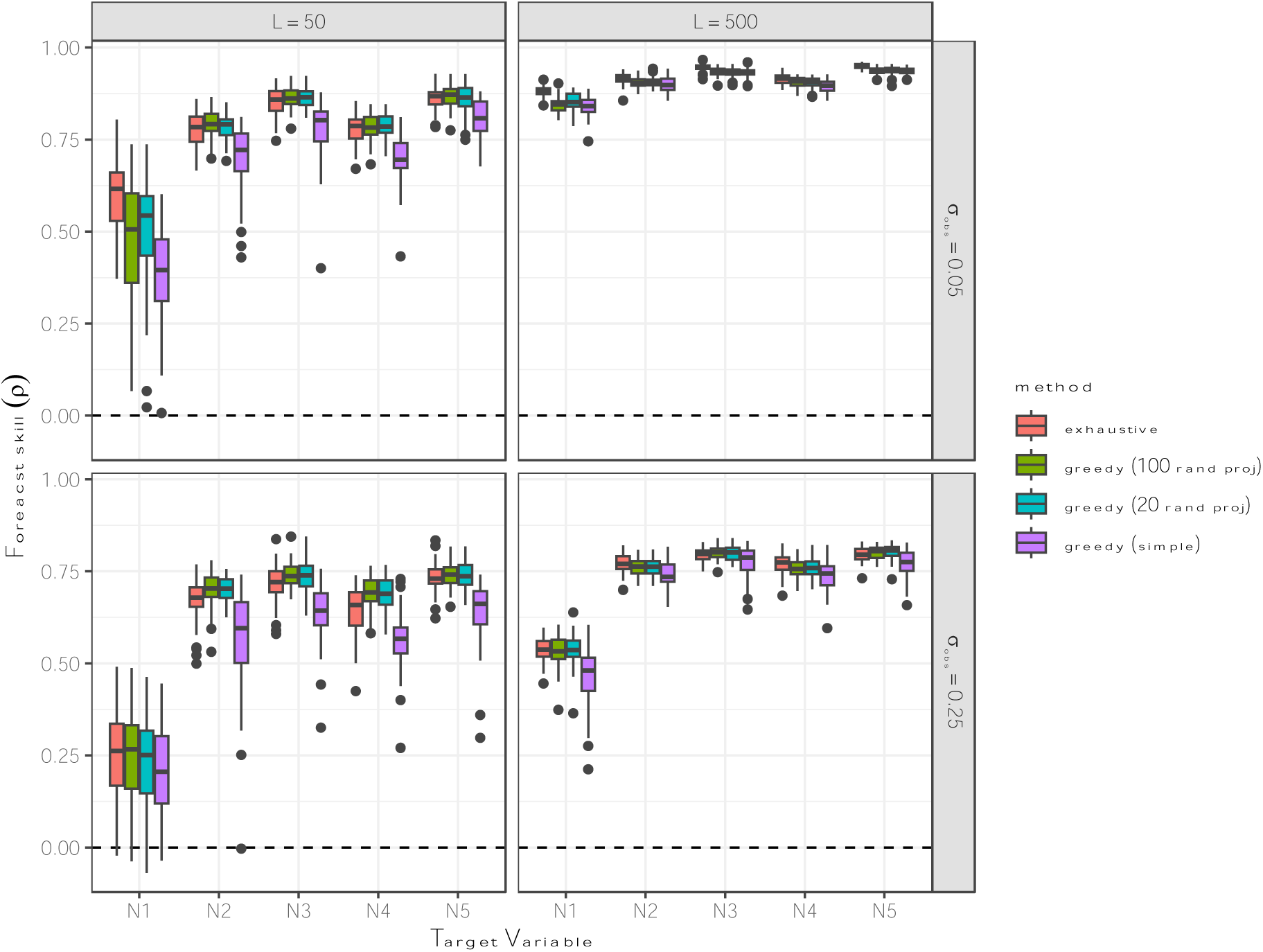
Out-of-sample forecast skill for multivariate EDM models constructed using four alternative algorithms applied to each population variable in the multispecies competition model. The algorithms included brute-force exhaustive search (red), greedy search with randomized projection coordinates and N_r_ = 100 re-randomizations (green), greedy search with randomized projection coordinates and N_r_ = 20 re-randomizations (teal), and greedy search using standard multivariate embeddings without random coordinates (magenta). Each model construction method was tested against the same set of randomly initialized model runs. The out-of-sample forecast skill for each method across tests are shown as standard bar-and-whisker plots in a 2x2 matrix of subplots. The subplot row corresponds to varying the degree of observational noise (σ_obs_) and column corresponds to the number of library points (*L*), corresponding to subplot column.

Using 100 re-randomizations of the projection and a maximum embedding dimension of 6 requires at most 10,500 multivariate forecasts. If more variables or more coordinates need to be considered, the number grows slower than the product of *k*, *m*, and *E*. Adding another 10 variables would require 16,500 simplex calculations.

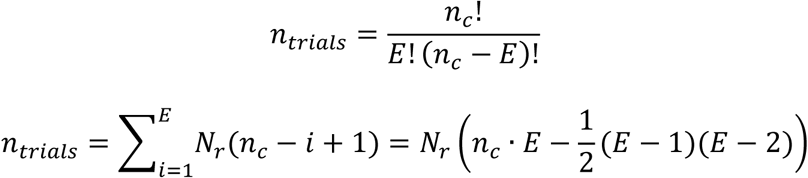

The most consequential feature of the experimental results (Fig 5 and S6), then, is that the out-of-sample forecast skill of models selected with the greedy random coordinate search was nearly comparable when only 20 re-randomizations were used instead of 100.

## 4 Conclusion

Reassuringly, the particulars of the algorithm do not seem critical to the immediate usefulness of the general concept. Randomized embeddings do appear to offer greater efficiency than Multiview-inspired random variable selection. From the present analyses, it is not clear if there is a benefit to identifying drivers by removing variable, replacing variable with randomized surrogate, or (not explicitly addressed here) replacing with another random projection coordinate. It may be that considering a sufficiently broad set of conditions for application there will be identifiable cases where each possibility is most strategic. A non-exhaustive list includes: total number of observations, observation error, process error (stochastic or high-dimensional forcing), missing variables, and confounding variables. Additionally, Fig S2 suggests that the algorithm is robust to both over and under embedding (using an embedding dimension lower or higher than necessary), but the individual algorithmic designs may be more or less so.

At this point many readers will have also noticed we’ve relied exclusively on greedy search for system identification which has been well known for quite some time to be, shall we say, suboptimal. Taking the ingredient of random projection embeddings, all the options for moving beyond greedy search lay ahead. Rather the demonstrations above should reinforce that there is abundant territory for applying empirical dynamic modeling where traditional time series length may be much more restricted than in noteworthy studies in the past, but overall data set size is orders of magnitude larger than pure time-series applications of EDM (8).

**TABLE 1.** Symbols and abbreviations for meta-parameters of numerical tests.

| SYMBOL | DESCRIPTION |
| --- | --- |
| $n_c$ | Number of candidate/covariate time series. |
| $N_r$ | Number of coordinate randomizations performed for a multivariate EDM test. |
| T:S | Ratio of true interacting variables to spurious candidate variables in numerical experiment. |
| L | Size of the library, i.e. number of vectors in the training set. |

## Acknowledgements

This work was supported by funding from the DoD-Strategic Environmental Research and Development Program 15 RC-2509, NSF DEB-1655203, NSF ABI-1667584, DOI-NPS-P20AC00527, and NSF OCE-2209284.

## Author Contributions

ERD, GP, and GS conceived the ideas and designed methodology; ERD performed and analyzed the numerical experiments; ERD led the writing of the manuscript. All authors contributed critically to the drafts and gave final approval for publication.

## Data Availability

Complete code to recreate the figures and results contained with this manuscript are provided in a Github repository that will be made public at the time of publication.

---

**Fig S1.**
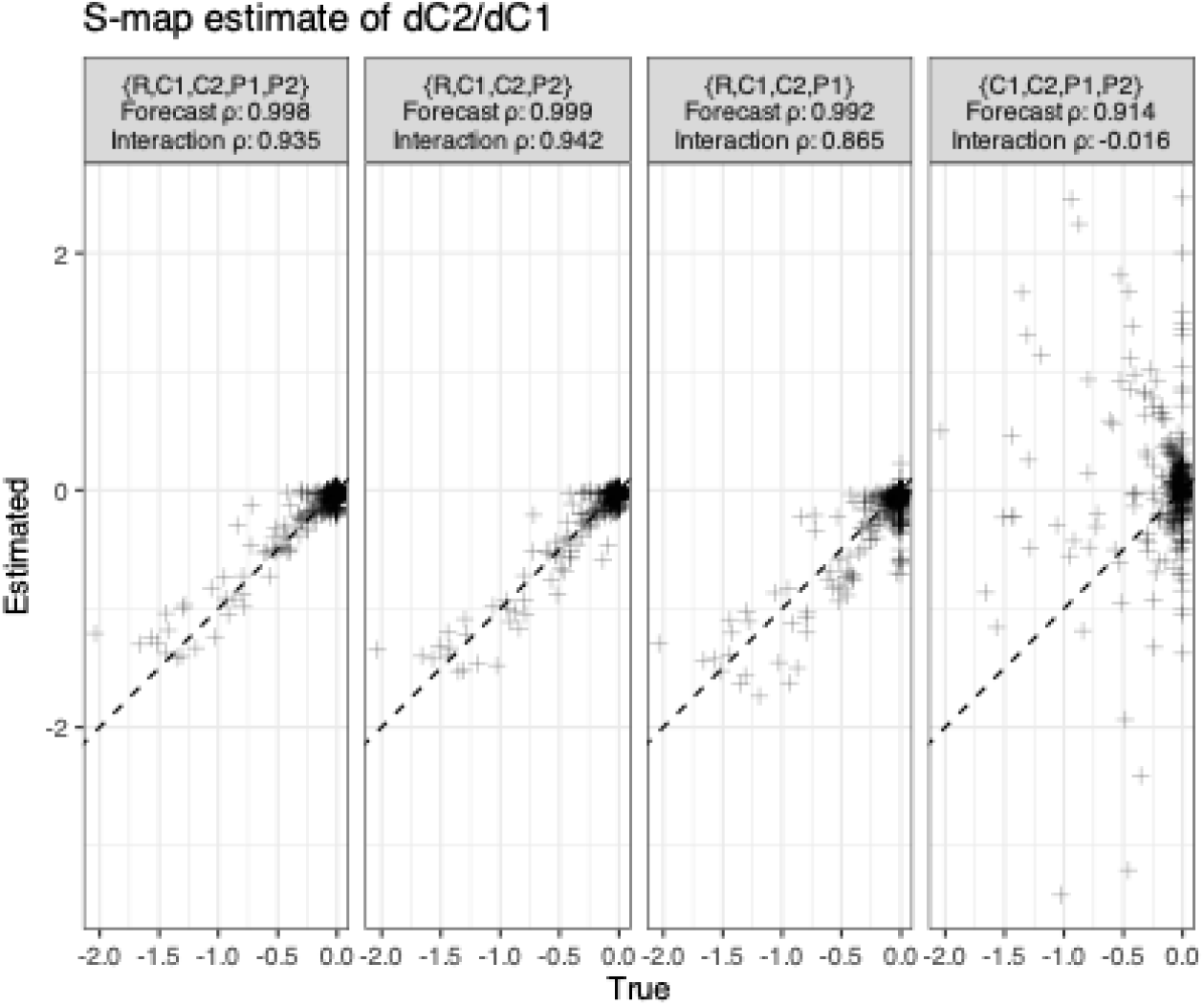
Differential sensitivity of interaction estimation to missing variables in a coupled food-chain model. While the realized competitive interaction can be reasonably estimated with S-map whether or not they include higher trophic levels (“full”, “no P1”, “no P2”), estimates of this interaction when the shared resource time series is missing are often erroneously estimated as positive. The S-map forecasts with that embedding are still largely accurate (ρ=0.891), despite the local linear model coefficients bearing little similarity to the true model (ρ=0.064).

**Fig S2.**
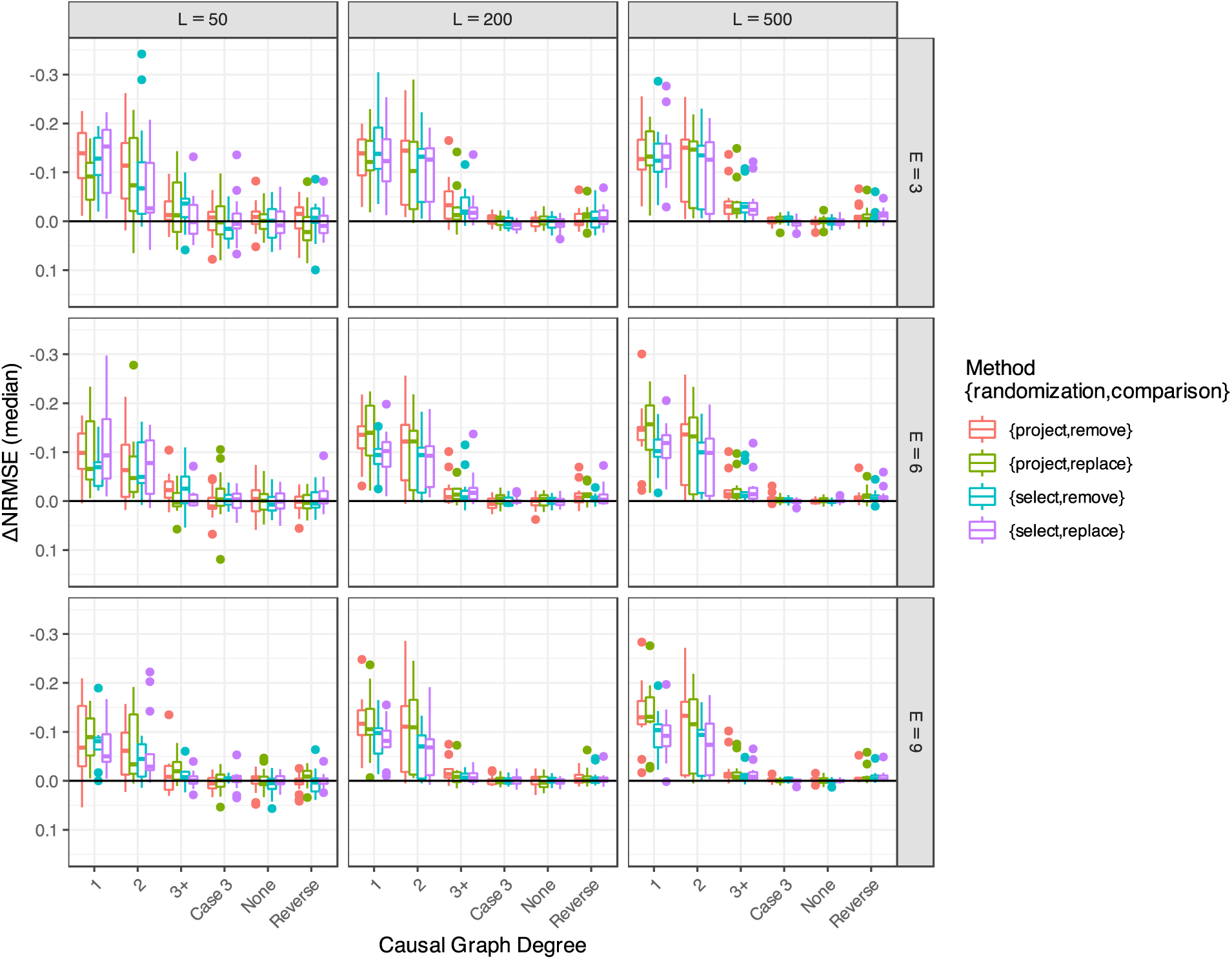
Comparison of different implementations of the general test. The same tests were run varying the design parameters of the analysis, including how many similar, but extraneous variables were included as possible candidates (row of plots) and the number of coordinates used for EDM forecasting (column of plots), i.e. the embedding dimension, *E*.

**Fig S3.**
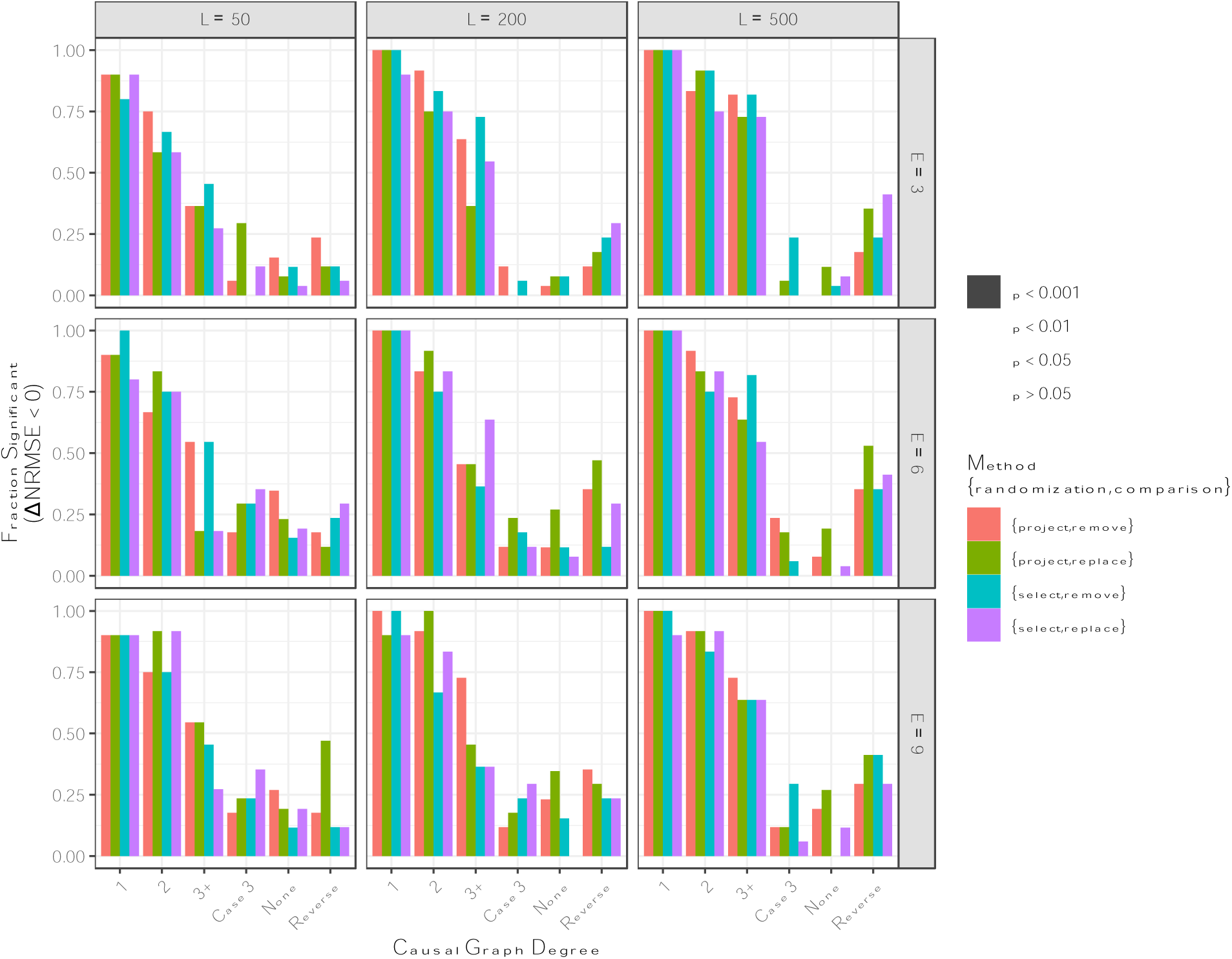
Comparison of different implementations of the general multivariate test via Wilcoxon test. The same tests were run varying the design parameters of the analysis, including the number of vectors in the training library (columns of plots) and the number of coordinates used for EDM forecasting (rows of plots), i.e. the embedding dimension, *E*. The filled bars indicate the fraction of variables that showed a significant increase in forecast skill (ΔNRMSE < 0), with lighter fill signifying progressively a weaker threshold for significance (*p* < 0.001, 0.01, and 0.05).

**Fig S4.**
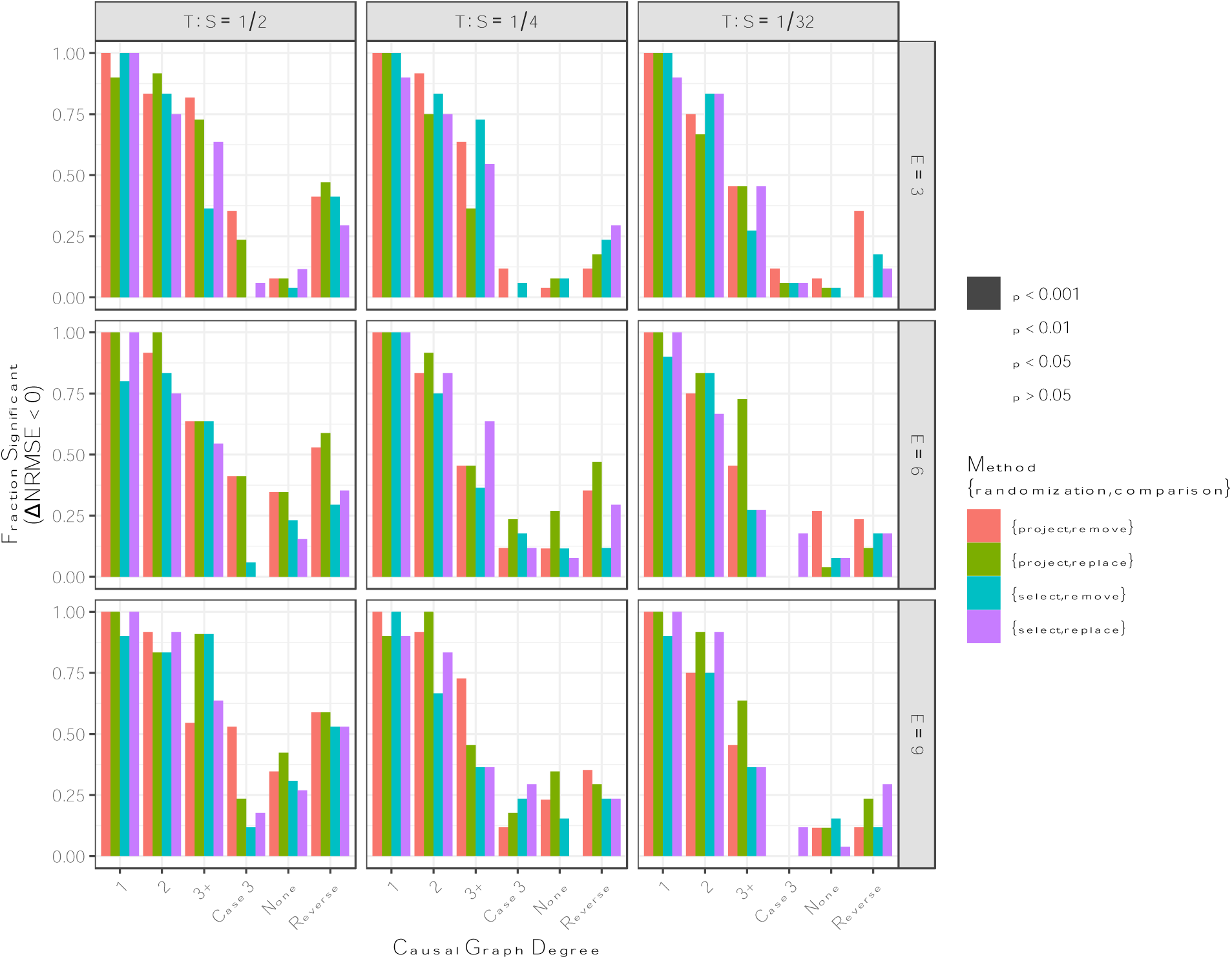
Comparison of different implementations of the general multivariate test via Wilcoxon test. The same tests were run varying the design parameters of the analysis, including how many similar, but extraneous variables were included as possible candidates (columns of plots) and the number of coordinates used for EDM forecasting (rows of plots), i.e. the embedding dimension, E. The filled bars indicate the fraction of variables that showed a significant increase in forecast skill (ΔNRMSE < 0), with lighter fill signifying progressively a weaker threshold for significance (*p* < 0.001, 0.01, and 0.05).

**Fig S5.**
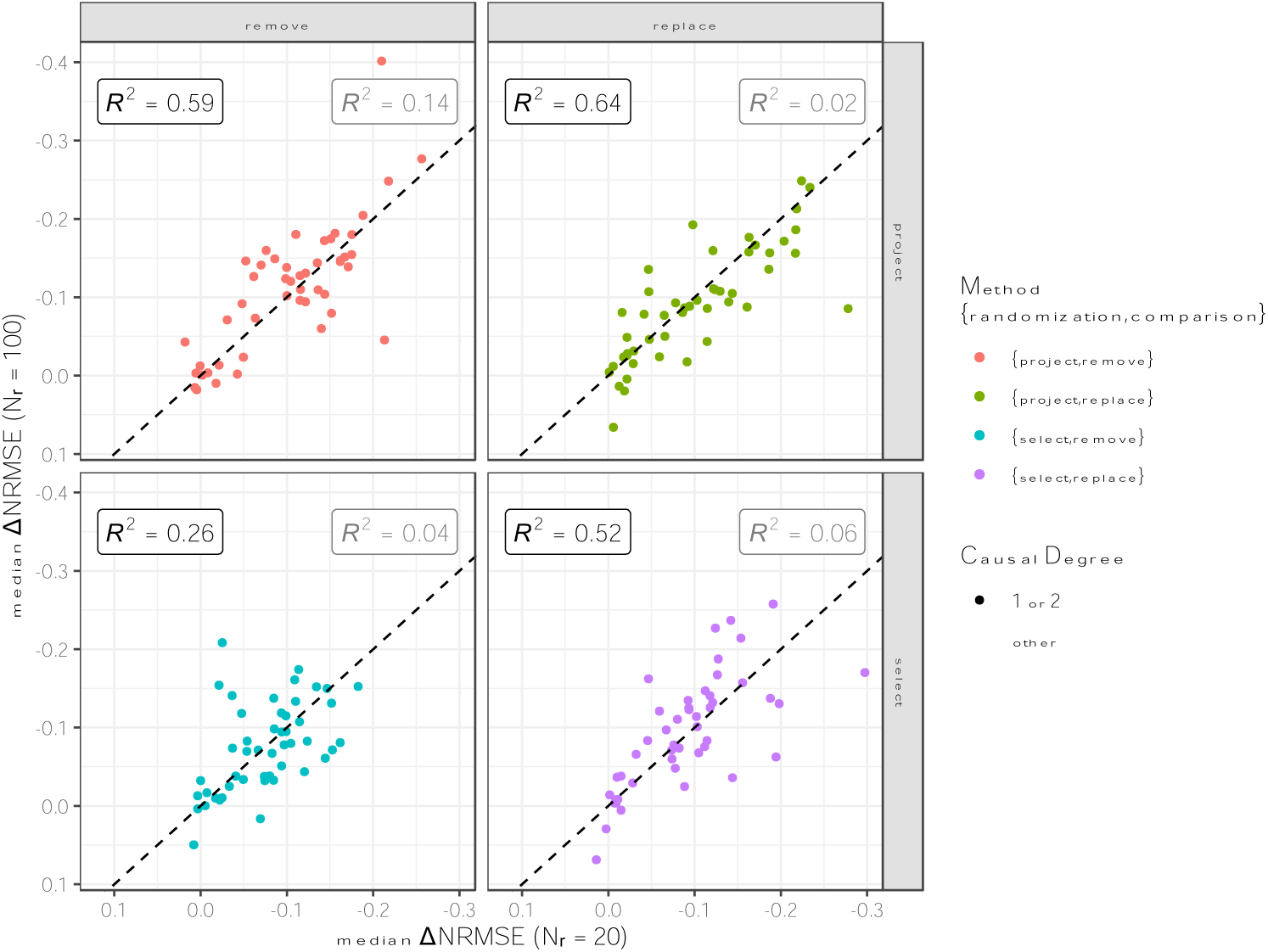
Recovering representative measure of forecast improvement (ΔNRMSE) using fewer re-randomizations (L = 200, T:S = 1:3, and E = 6). The correspondence between ΔNRMSE calculated for N_r_=100 and N_r_=20 is shown between different methods of randomization (randomly projected coordinates, randomly selected coordinates) and comparison (remove, replace). R^2^ values are plotted for tests between closely connected variables (causal degree ≤ 2) in black on the left; R^2^ values for the other remaining variable pairs are shown in grey on the right.

**Fig S6.**
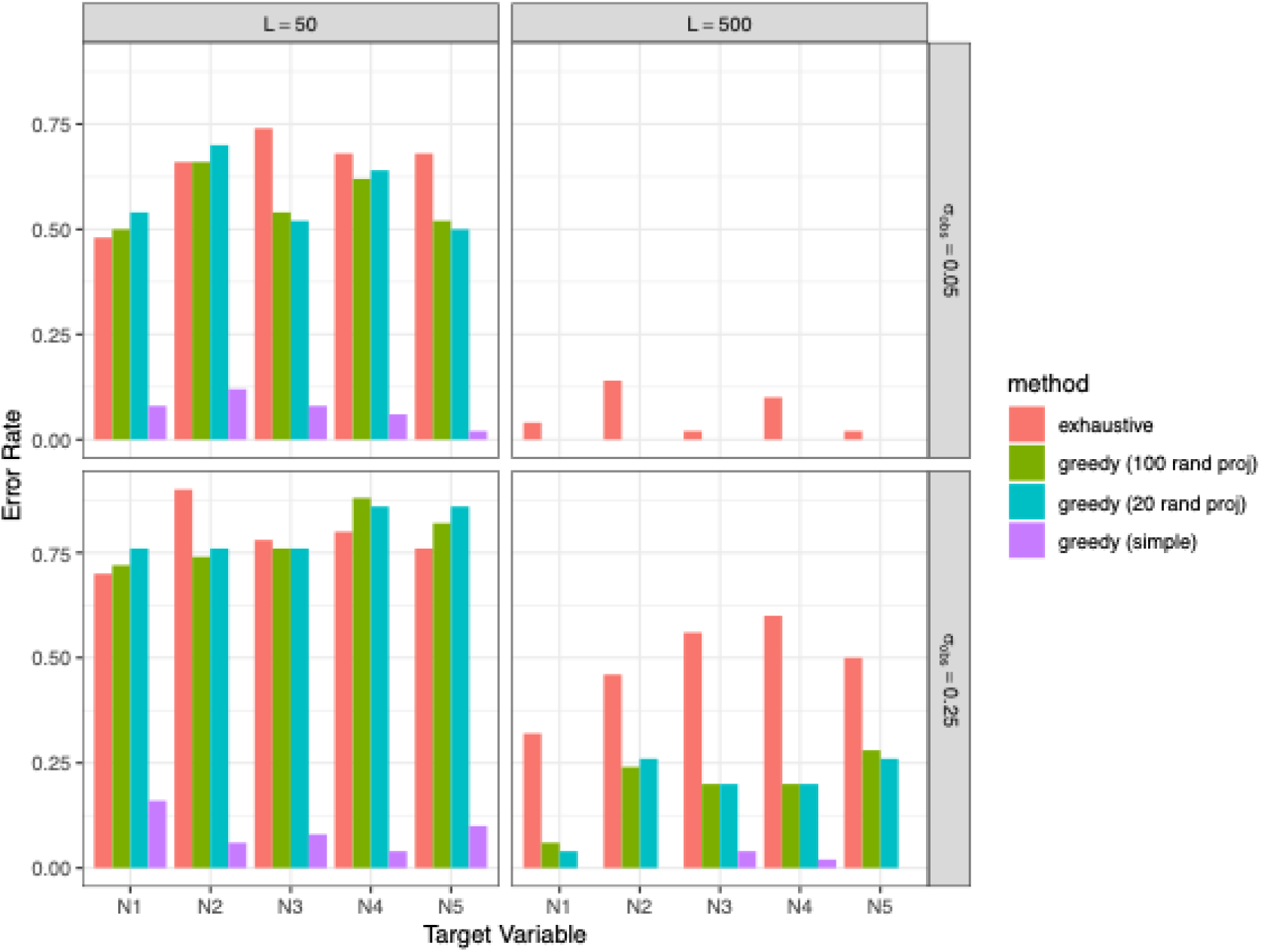
Error rate of selecting at least one erroneous time-series variable by different model selection methods in experiment shown in Fig 6. The algorithms included brute-force exhaustive search (red), greedy search with randomized projection coordinates and a resample of 100 (green), greedy search with randomized projection coordinates and a resample of 20 (teal), and greedy search using standard multivariate embeddings without random coordinates (magenta).

**Table S1.** Parameter values for Repressilator network model described in Fig 3. Note that the Hill coefficient representing the shape of saturating binding dynamics is labeled here with *h* rather than *n* (as in (27)) to avoid confusion with other uses of *n* as a symbol.

|  |  |
| --- | --- |
| $\alpha_D$ | 35 |
| $\beta_D$ | 2 |
| $\kappa_D$ | 45 |
| $ks0_D$ | 1.0 |
| $ks1_D$ | 0.025 |
| $h_D$ | 2.6 |
| $\eta_D$ | 2.0 |
| $\alpha_{R1}$ | 25 |
| $\beta_{R1}$ | 1 |
| $\kappa_{R1}$ | 40 |
| $ks0_{R1}$ | 1.0 |
| $ks1_{R1}$ | 0.025 |
| $h_{R1}$ | 2.6 |
| $\eta_{R1}$ | 2.0 |
| $\gamma_{R1}$ | 0.3 |
| $\alpha_{R2}$ | 32 |
| $\beta_{R2}$ | 2 |
| $\kappa_{R2}$ | 40 |
| $ks0_{R2}$ | 3.0 |
| $ks1_{R2}$ | 0.027 |
| $h_{R2}$ | 2.65 |
| $\eta_{R2}$ | 0 |
| $\gamma_{R2}$ | 0.2 |

## Notes

### Competing Interest Statement

The authors have declared no competing interest.

